# Dynamic gait transition in the *Scolopendromorpha scolopocryptops rubiginosus L. Koch* centipede

**DOI:** 10.1101/312280

**Authors:** Shigeru Kuroda, Nariya Uchida, Toshiyuki Nakagaki

## Abstract

Crawling using locomotory waves is a common method of locomotion for limbless and many-legged invertebrates. It is generally believed that the direction of locomotory waves is fixed for a given species. However, by recording and performing detailed analyses of the gait patterns of a *Scolopendromorpha scolopocryptops rubiginosus L. Koch* centipede in various conditions, we found that it dynamically generated its gait to allow for locomotory waves that varied in direction. By introducing the *wave-index* order parameter to characterise locomotory waves, we showed that gait patterns were associated with control of stride rather than rotation frequency.

## I. INTRODUCTION

Biological locomotion in uncertain environments is a highly adaptive behaviour that is achieved by dynamic gait generation. Crawling is a fundamental method of locomotion for both limbless and legged invertebrates. Gait patterns are often observed as waves propagating along the major body axis, such as peristaltic waves in Annelida such as earthworms [1], pedal waves in Mollusca such as snails [2–5], and leg-density waves in Myriapoda such as millipedes and centipedes [6, 7]. Myriapoda locomotion is usually accompanied by density waves formed by leg-tip positions along the body axis. These waves are referred to as *locomotory waves* in this paper. Although the propulsive machinery is very different among these crawlers, it has been shown that the locomotory waves are kinematically similar to each other [8] and play the common mechanical role of pushing the body forward [9, 10].

Locomotory waves are classified according to their propagating direction [11]. They are called *direct* if they propagate in the movement direction, and *retrograde* if they propagate in the opposite direction. The direction of locomotory waves for a given species is commonly considered fixed. For example, polycaeta worms, snails, and millipedes use direct waves, while earthworms, chiton, and one kind of centipede use retrograde waves [8]. We refer to this idea as the *one-species-one-wave hypothesis*. Centipedes (class Chilopoda) are soft-bodied arthropods with many legs. There are five orders of centipedes, and their gaits have been characterised by order [7]. Scutigeromorpha and lithobiomorpha use direct waves while Scolopendromorpha and others use retrograde waves. Although it is believed that differences in body features and habitat may cause the order-dependency of the locomotory wave direction, the determining factor for the selection of the wave direction is poorly understood.

In natural circumstances, crawlers migrate under various conditions, and should therefore possess some ability to adapt their gait. Laboratory observations about gait-modulation of crawlers have been made [9, 12–15]. Manton [8]showed that the gait patterns of many-legged organisms are characterised by wavenumber (the phase difference between successive legs), rotation frequency (the multiplicative inverse of the duration of a cycle of a leg movement), duty factor (the relative duration of the recovery and power strokes by the legs), span (the moving distance during a power stroke of a leg), and stride (the moving distance during a cycle of a leg movement, i.e. stride = span / duty factor). The organisms can speed up by varying leg movements, including increasing the rotation frequency, increasing the span, and decreasing the duty factor. Each species weighs the importance of each method differently.

We examined the gait patterns of the Scolopendromorpha species of centipede under different conditions and found they possess various modes of locomotion that use retrograde waves, direct waves, and a combination of both that we will call *splitting waves*. That is, a counterexample of the one-species-one-wave hypothesis was discovered. By introducing an order parameter called the *wave-index*, we characterised the locomotory waves and showed that gait patterns are associated with control of stride rather than rotation frequency. Finally, we discussed the adaptive locomotion of invertebrate crawlers.

## II. ORGANISMS AND METHODS

### A. Animals

The experiments were performed using centipedes *(Scolopendromorpha scolopocryptops rubiginosus L. Koch*) (*n* = 16; body length= 46.4±6.8 mm, weight= 0.28*±*0.04 g (mean *±* std); all were putative adults) collected in Hakodate, Japan. These centipedes have 23 paired legs, and each leg is referred to by a number as shown in Fig. 1a. The foremost and hindmost leg pairs are not walking legs; they are used for feeding and stabilisation during locomotion, respectively. Eighteen pairs of walking legs (the 3rd through the 20th) were analysed in this paper. Three pairs near the edges (pairs 2, 21, and 22) were excluded because their movements were not large enough for analysis. Gait patterns of the millipede *Polydesinida oxidus* (*n* = 4) were also observed for comparison. This millipede has 20 segments and 31 legs per side. The 2nd through 4th segments (called haplosegments) have a single pair of legs each, and the 5th through 18th segments (diplosegments) have two pairs of legs each. The legs are numbered from the head to tail in increasing order. All experiments were performed at room temperature (23±2 °C). We observed their locomotion in various situations.

**FIG. 1.**
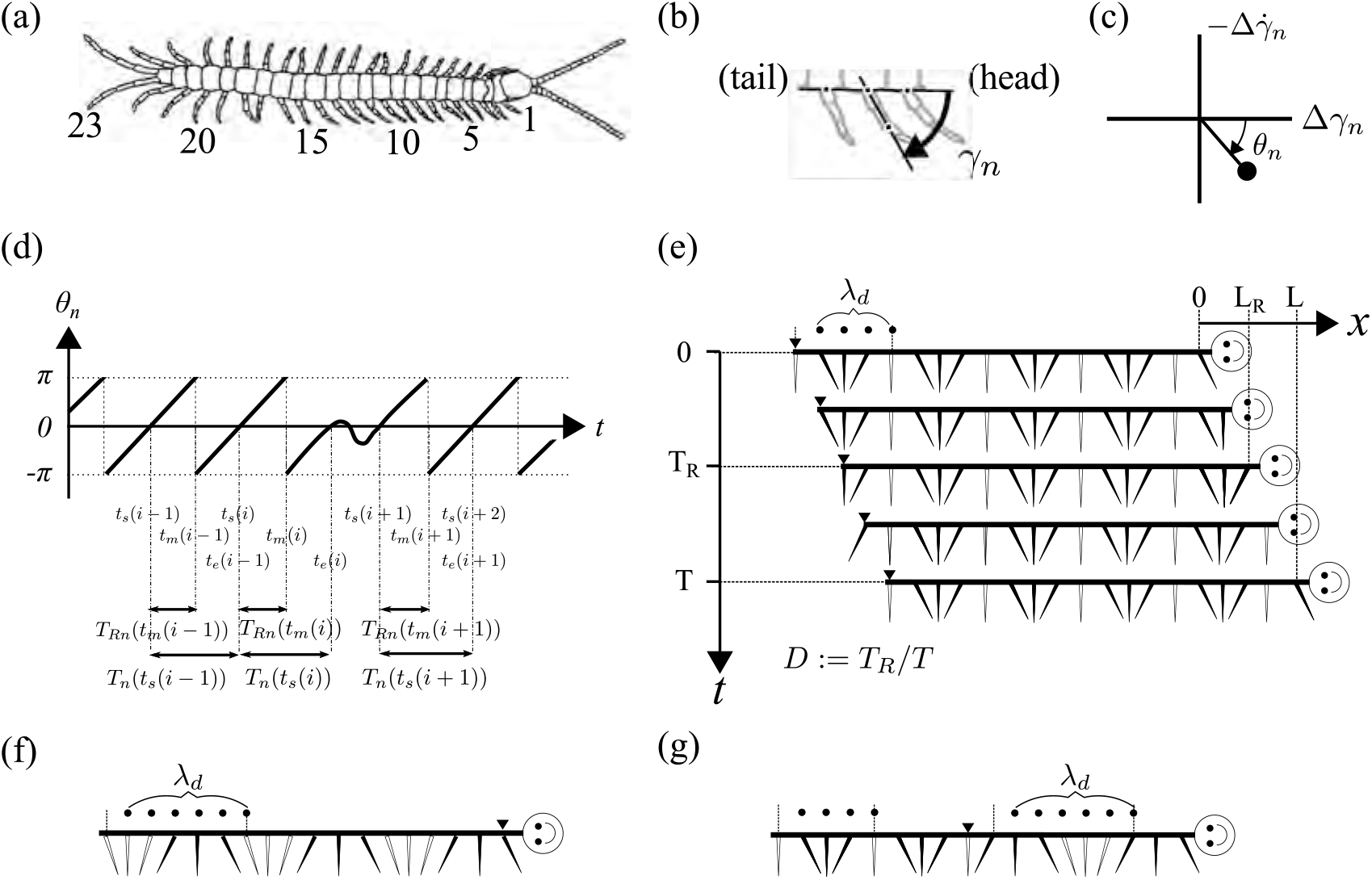
(a) Top view of a centipede showing the numbering of the 23 paired legs. (b) Swing angle of the *n*th right leg, *γ_n_*. (c) Phase of the *n*th leg, *θ_n_*. (d) Retraction period *T_n_*(*t_s_*(*i*)) = *t_m_*(*i*) – *t_s_*(*i*) and the pace duration *T_n_*(*t_s_*(*i*)). (e) Diagram of a gait with retrograde waves showing the dominant wavelength λ_*d*_ [#leg], retraction period *T_R_*, pace duration *T*, span *L_R_*, stride *L*, and duty factor *D*. Black filled lines and blank lines denote the legs during the retraction and protraction periods, respectively. (f) Diagram of a snapshot of a gait with direct waves. (g) Diagram of a snapshot of a gait with split waves. The black triangles in (d-f) show the most phase-advanced legs in each gait.

### B. Apparatus

#### 1. Walking and running on a flat surface experiment

In the walking or running on a flat surface experiments, the animals crawled in lanes on a plastic tray. Wet paper (Nippon Paper Crecia, Kimwipe s-200) or sandpaper was used to change the floor surface condition. Lightly tapping the tail of the centipede induced high-speed movement (< 120 mm/s), while non-stimulated centipedes performed exploratory crawling at relatively low speeds (< 30 mm/s).

#### 2. Swimming experiment

In the swimming experiments, a pool about 20 mm deep was prepared and the animals were gently placed on the surface of the water. Three pools each filled with pure water, 1.5w/v% and 2.0w/v% carboxymethylcellulose (CMC) solutions were prepared.

#### 3. Slippery slope experiment

In the slippery slope experiments, a narrow plastic tray (13 mm) with high walls (60 mm) was tilted at a desired angle (0–30°).

### C. Measurements

Images were taken with a digital camera (EX-F1, Casio) at 300 frames per second. Animal movements were recorded from above or from the side. Videos taken from above were primarily used for analysis; those from the side were for supplementary use.

### D. Data analysis

Colour movies were analysed after conversion to greyscale. Image brightness and contrast were optimised based on analysis of image histograms. Image analyses were performed using pre-installed and custom-made plug-in programs for the ImageJ software [16].

Space–time plots like those in Fig. 2 were used to visualise and roughly inspect the development of the wave pattern over time. A suitable one-dimensional region of interest (ROI), which is the segmented line in the plot, was chosen to demonstrate the evolution of waves on that side. In cases where the crawler changed posture such that the ROI did not cover all legs, we redefined the ROI to fit the new posture within the frame. When doing so, the previous ROI was smoothly transformed into the new ROI over several dozen frames.

**FIG. 2.**
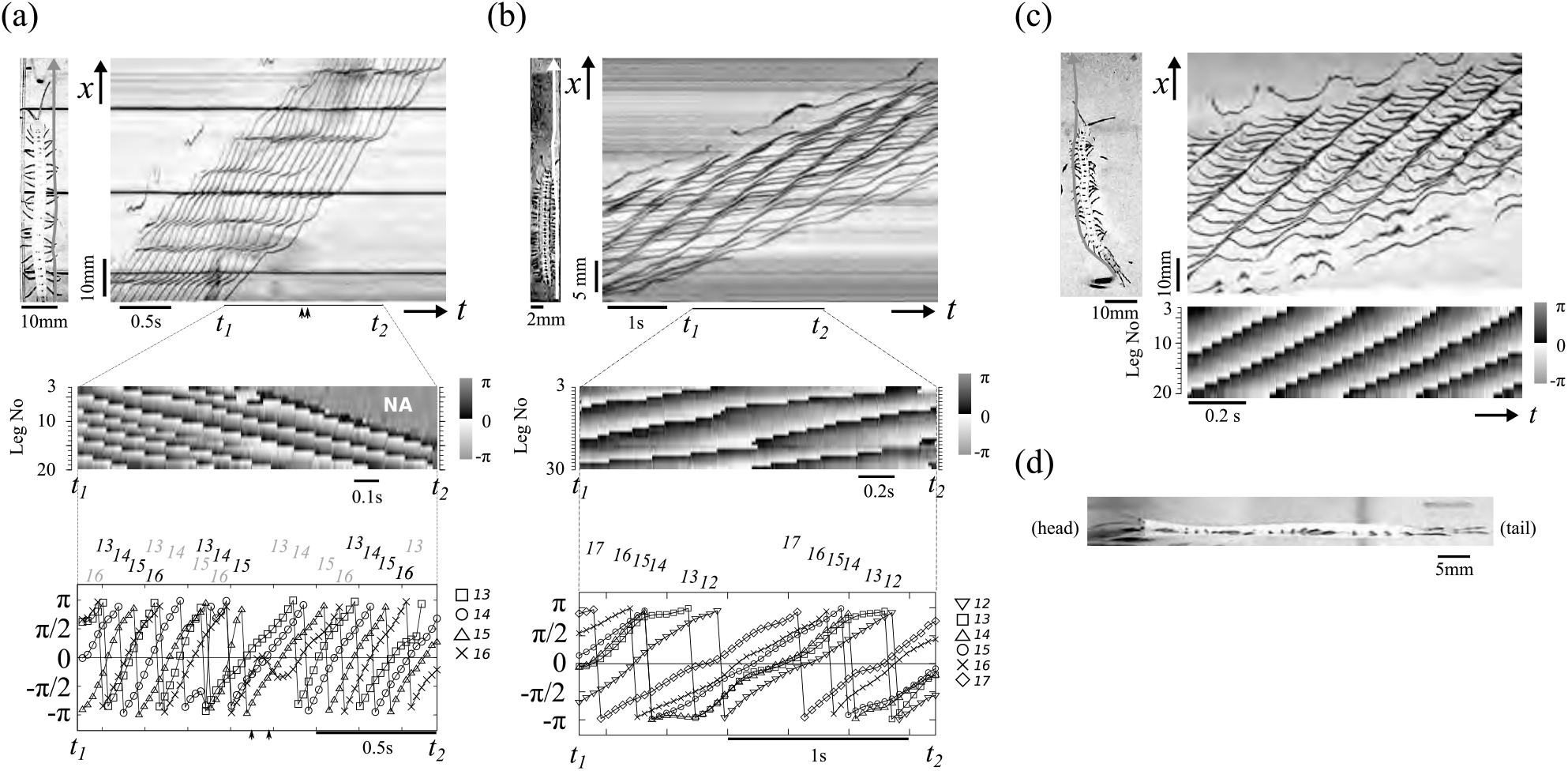
(a) Retrograde waves of a centipede during free walking. The upper-right panel shows a space–time plot, in which each stripe shows the trajectory of each leg’s position on the one-dimensional axis shown in the upper-left panel. The middle panel shows a phase map of the 3rd through 20th legs between *t*_1_ and *t*_2_. The bottom panel shows time courses of the phase of four successive legs between *t*_1_ and *t*_2_. The leg number is shown above the graph where the leg phase reaches *π*, which corresponds with the end of the power stroke. (b) A space-time plot (upper panel) and a phase map (bottom panel) of direct waves of a millipede during free walking. The phase map of 24 legs (from the 3rd to the 30th legs; legs 9, 24, 35, and 29 are missing) is shown between *t*_1_ and *t*_2_. The bottom panel shows the time courses of the phase development of successive legs (from the 12th to the 17th legs) between *t*_1_ and *t*_2_. Leg numbers were assigned in the same way as in (a). (c) A space–time plot (upper panel) and a phase map (bottom panel) of direct waves in a centipede during water crawling. (d) A side view of the centipede. The head is on the left.

To quantify each leg’s movement relative to the body and other legs, several characteristics were estimated as follows:

#### 1. Swing angle and cycle phase

The positions of the base and a point between the middle and tip of each leg were tracked. The swing angle of the leg was determined as the angle between a line through the base and the leg midpoint and a line through the bases of the legs on either side (see Fig. 1b). The cycle phase of the *n*th leg’s movement relative to the body (*θ_n_* ∈ [−*π*, *π*]) was determined from the displacement of the swing angle (*γ_n_*) from its average over time 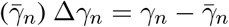 and its time derivative 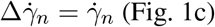 (Fig. 1c).

#### 2. Retraction period, pace duration, rotation frequency, and duty factor

After obtaining the time series of the phase of the *n*th leg ({*θ_n_*(*t*)}), the *i*th cycle of the leg movement was determined as the duration from *t_s_*(*i*) to *t_e_*(*i*), where *t_s_*(*i*) and *t_e_*(*i*) are consecutive times that satisfy the following conditions: *θ_n_*(*t_s_*(*i*)) = *θ_n_*(*t_e_*(*i*)) = 0 and *θ_n_*(*t*) ≠ 0 for ∀*t* ∈ (*t_s_*(*i*),*t_e_*(*i*)), and ∃*t*\* ∈ (*t_s_*(*i*), *t_e_*(*i*)) such that *θ_n_*(*t*\*) > *π*/4. The last condition was adopted to exclude the duration of tentative motions before the retraction.

The *i*th cycle was divided into a *retraction period* [*t_s_*(*i*),*t_m_*(*i*)] and a *protraction period* [*t_m_*(*i*),*t_e_*(*i*)], where *t_m_*(*i*) ∈ (*t_s_*(*i*),*t_e_*(*i*)) is the earliest time at which the phase becomes *π*. The period of the cycle (the *pace duration*) (*T_n_*(*t_s_*(*i*))) and the retraction period (*T_Rn_*(*t_m_*(*i*))) were estimated as *T_n_*(*t_s_*(*i*)) = *t_e_*(*i*) – *t_s_*(*i*) and *T_Rn_*(*t_m_*(*i*)) = *t_m_*(*i*) – *t_s_* (*i*), respectively. The *duty factor* (*D_n_*(*t_m_*(*i*))), which is the ratio of the retracting period over a cycle, was estimated as *T_Rn_*(*t_m_*(*i*))/*T_n_*(*t_s_*(*i*)) (Fig. 1e). Finally, the time series for the periods *T_n_*(*t*), *T_Rn_*(*t*), and *D_n_*(*t*) were reconstructed by linear interpolation of *T_n_*(*t_s_*(*i*)), *T_Rn_*(*t_m_*(*i*)), and *D_n_*(*t_m_*(*i*)), respectively, along the time axis. The rotation frequency *ω_n_* was estimated as the multiplicative inverse of the pace duration *T_n_*.

In this paper, for each set of time series, {*Z_n_*(*t*)}_*n*_, where *Z_n_* = *T_n_*, *T_Rn_*, *D_n_*, or *ω_n_*, the leg-averaged time series is denoted by *Z*(*t*).

#### 3. Phase relationship between adjacent legs, local and global waves, and the wave-index

The phase relationship between adjacent legs was defined in a standard way. If *θ_n_*(*t*) – *θ*_*n*+1_(*t*) < *π* (or > *π*) (mod 2*π*), the phase of the *n*th leg is ahead of or behind the phase of the *n* + 1th leg at *t*, and the two legs have an *antero-posterior* or *postero-anterior* phase relationship, respectively. If any two adjacent legs between the *n* + 1th and *n* + *q*th legs (*q* ≥ 2) have an antero-posterior phase relationship, the legs form a (*q-length*) *local antero-posterior wave*. The *n* + 1th leg or *n* + *q*th leg is called the *most phase-advanced* or *phase-behind* among these legs, respectively. Similarly, a *local postero-anterior wave* was defined. If a *q*_1_-length local antero-posterior wave and a *q*_2_-length local postero-anterior wave shared a most phase-advanced or phase-behind leg, the shared leg was called a *source* or *sink* leg, respectively, and the combined (*q*_1_ + *q*_2_ – 1)-length wave was called a *local splitting wave*. If there was a unique local antero-posterior wave including both the foremost and the hindmost walking legs, the wave was called a (*global*) *antero-posterior wave. (Global) postero-anterior waves* and *(global) splitting waves* were similarly defined (Fig. 1e-g). During forward locomotion, in particular, antero-posterior and postero-anterior waves were *retrograde* and *direct* waves, respectively. In this paper, we only investigated forward locomotion or swimming. Therefore, we rephrased the terms antero-posterior and postero-anterior as retrograde and direct, respectively, in this paper.

To characterise the mean phase relationship of legs along the body axis, we introduce the *wave-index*. The *wave-index* at time *t*, *I_w_* (*t*) ∈ [0, 1], was defined as the proportion of the number of legs in a postero-anterior relationship with the immediately following leg at *t*. The wave-index of a splitting wave with a source takes an intermediate value.

#### 4. Dominant (absolute) wavenumber and mean (signed) wavenumber

To robustly estimate the instantaneous wavelength of a phase wave along the body axis, we calculated the mean wavelength from local waves that persist over more than one wavelength (more exactly, 2*π* in terms of the phase). The dominant instantaneous wavelength, *λ_d_*, was estimated as the mean value of the instantaneous wavelengths during a mean period of leg movement cycling (Fig. 1e-g). The dominant (absolute) wavenumber in the phase wave, *κ_d_*, was estimated as 2*π*/*λ_d_*. We also estimated dominant wavenumbers in the local retrograde and local direct waves (*κ_d_*|_*retro*_, *κ_d_*|_*direct*_), respectively. The mean (signed) wavenumber, *κ_m_*, was estimated as the leg-averaged value of the signed phase difference (∈ [–*π*, *π*]) of each leg from its immediately following leg, i.e., *θ_n_*(*t*) – *θ*_*n*+1_(*t*) ∈ [–*π*,*π*]. The mean wavelength in the phase wave was estimated as 2*π*/*κ_m_*.

#### 5. Span, stride, and locomotion speed

A two-dimensional trajectory (*x*(*t*), *y*(*t*)) of a representative point on the crawler was obtained from the top-view video. The moving distance during a period (*t_i_*, *t_j_*) denoted as *ℓ*(*t_i_*, *t_j_*) was estimated by the length of the trajectory during the period. The span at time *t*, *L_R_*(*t*), was then estimated as the moving distance during the leg-averaged retraction period at that time, *L_R_*(*t*) = *ℓ*(*t* – *T_R_*(*t*)/2, *t* + *T_R_*(*t*)/2). Similarly, the stride at time *t*, *L*(*t*), was estimated as the moving distance during the leg-averaged pace duration at that time, *L*(*t*) = *ℓ*(*t* – *T*(*t*)/2,*t* + *T*(*t*)/2). The locomotion speed at time *t*, *V*(*t*), was estimated by *L*(*t*)/*T*(*t*). In the slippery-slope condition, we set the *x*-axis in the slope direction and estimated the span, stride, and velocity in that direction as *L*_*R*|*x*_(*t*) = *x*(*t* + *T_R_*(*t*)/2) – *x*(*t* + *T_R_*(*t*)/2), *L*|*x*(*t*) = *x*(*t* + *T*(*t*)/2) – *x*(*t* – *T*(*t*)/2), and *V*|_*x*_ = *L*|_*x*_(*t*)/*T*(*t*), respectively.

#### 6. Leg-tip position

In Section IIIC, the putative leg-tip position of the *n*th leg, *X_n_*(*t*), along the body axis was estimated as *b_n_* + *r_n_cos*(*θ_n_*(*t*)) (*n* = 3,…, 18), where *b_n_* is the coordinate of the base of the nth leg along the body axis and *r_n_* is the length of the nth leg. For simplicity, each *b_n_* was assumed constant throughout the locomotion.

When leg data were partially unobtainable, the characteristics were estimated from the other legs in Figs. 2a, 3, and 5a. However, in Fig. 5, they were estimated only when all the legs’ data were available, because the wave-index and wavenumber are sensitive to a lack of data.

**FIG. 3.**
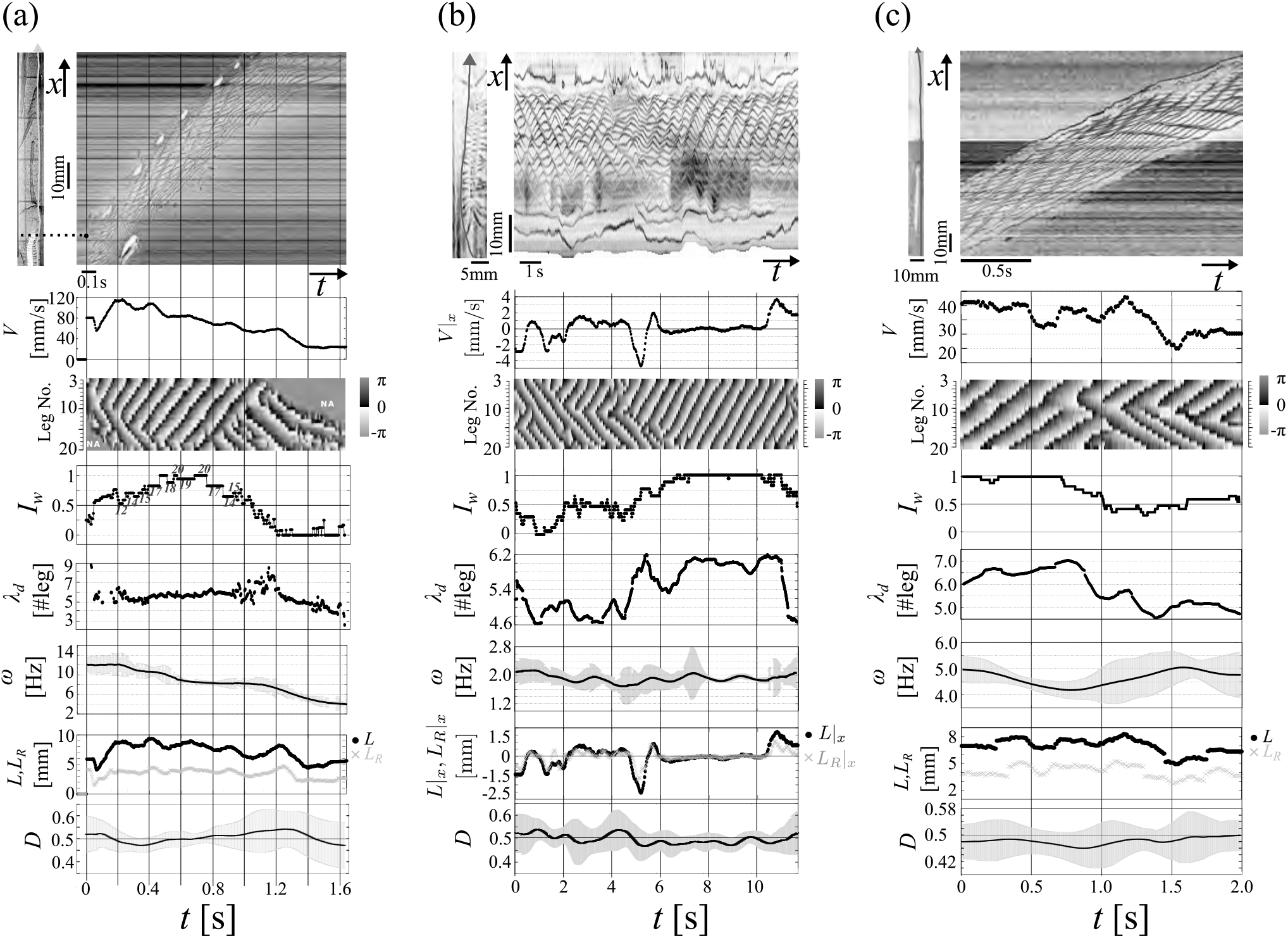
Continual transition in a locomotory wave under three conditions. For each condition, a space-time plot of the position of the legs on one side, the time course of speed (*V* or *V*|_*x*_), a phase map, the wave-index (*I_w_*), the wavelength (λ), the rotation frequency (*ω*), the stride (*L* or *L*|_*x*_), the span (*L_R_* or *L_R_*|_*x*_), and the duty factor (*D*) are vertically arranged with the same time axis. In the graph of the mean pace period and duty factor, the vertical grey bar shows the standard deviation of the cycle over the legs. (a) Gait in the dash and slowing conditions. The upper-left panel shows the spatial axis at the first frame, and the black dot near the left edge in the upper-right panel shows the head position before the dash. In the wave-index graph, the global source leg numbers are indicated near where they appeared. In the phase map, “NA” indicates unavailable data. (b) Gait change of a centipede on a 25° slippery slope. (c) Gait change of a centipede transitioning from wet paper to plastic.

## III. EXPERIMENTAL RESULTS

### A. Stable gait (1): Free walking on a solid flat surface

The upper panel in Fig. 2a shows a space-time plot of the leg-density waves of a centipede during free locomotion on a solid flat surface. Dense and sparse wave phases remained stationary with respect to the ground, but propagated from the head to the tail along the body axis. This is shown in the phase map (the middle panel in Fig. 2a). Therefore, the waves were retrograde waves, which propagate opposite to the direction of motion. The dense parts of retrograde waves consist of the legs that are nearly stationary with respect to the ground during power strokes, while the sparse parts consist of the legs that move forward with respect to the ground during recovery strokes [9]. A tentative motion before retraction in the 16th leg (the period between the black arrowheads) and lengthened protraction period in the 17th to 20th legs were observed. In contrast, the leg-density waves of a millipede were direct waves that propagated in the same direction as the motion (Fig. 2b). In the direct waves, the correspondence between wave phases and the stroke period was opposite to that in the retrograde case; that is, the sparse part of the wave consisted of legs nearly stationary relative to the ground in the power stroke. During the recovery stroke, the dense parts consisted of forward-moving legs.

### B. Stable gait (2): Swimming on the surface of the water

We used a water pool to observe crawling under different conditions from the flat solid substrate. After the animal was gently placed on the water, it proceeded to swim over the water after a brief struggle. Surprisingly, direct waves with large body undulations appeared during swimming (Fig. 2c). In this case, the dense parts of waves consisted of legs moving over the water surface performing the recovery stroke, and the sparse parts consisted of legs moving beneath the water surface performing the power stroke (Fig. 2d). The swimming with direct waves was also confirmed in the viscous CMC solutions. This suggests that direct waves are one stable mode for centipede locomotion, and that the direction of locomotory waves can change according to substrate conditions. However, further experiments indicated that changing surface conditions is not necessary to change the wave direction.

### C. Continual gait transition (1): Dashing and slowing down

When centipedes’ tails were lightly tapped, they sometimes ran (dashed) at high speed, and direct waves were often observed. Figure 3a shows a space–time plot of a centipede’s right-side legs after tapping its tail in which we observed a continual gait transition. Immediately after tapping the tail (*t* = 0), several consecutive anterior legs (1st to 11th, except the 3rd) simultaneously kicked the ground. This resulted in a sudden acceleration from 0 mm/s to 80 mm/s, starting from the time of the tap. During the successive recovery phases of these legs, a phase difference gradually formed between the legs. Two source legs also appeared along the body axis, where the anterior source was around the 6th leg and the posterior source may have been around the hindmost leg (this remains unconfirmed). As the phase developed, the anterior source shifted posteriorly, until the posterior source was eventually eliminated when the 12th leg become the global source (*t* ≈ 0.2 s). In this period, increased speed accompanied increased span and decreased duty factor, but the rotation frequency was constant. After reaching the maximum speed (< 120 mm/s), the speed slowly decreased until around *t* = 1.5 s, and this was accompanied by increased pace duration and duty factor, but the span was constant. During this period, the wave sources intermittently moved from the middle to the tail (wave-index *I_w_* = 0.5 ↗ 1), where the phase relationship between the source leg and its immediately posterior leg was reversed. The direct wave was maintained for a short time. Following that, the wave source gradually moved from the tail to the head (*I_w_* = 1 ↘ 0), where phase reversals tended to be allowed between non-source legs. During this period, the speed again gradually decreased with an increased duty factor and decreased span, and the rotation frequency was constant. After establishment of the retrograde waves, the speed sharply decreased (*t* = 1.2–1.5 s) with decreased rotation frequency and span and increased duty factor. Until *t* = 1.2 s, the dominant wavelength remained high (*λ_d_* > 5) compared with the dominant wavelength during free walking (*λ_d_* < 5).

To see how the legs controlled their mutual positions to realise the gait transition, we plot the time series of leg-tip positions *X_n_*(*t*) in Fig. 4. The leg-tip positions *X_n_*(*t*) for 3 ≤ *n* ≤ 18 are shown in Fig. 4(a). Wavefronts of propagating phase waves are indicated by dotted lines. The wavefronts with positive or negative gradients correspond to local direct or retrograde waves, respectively. To avoid curves overlapping each other, which makes it difficult to find the wavefronts, the intervals of adjacent bases were plotted wider than the actual intervals in Fig. 4(a) by shifting the origins of *X_n_* downwards by 2*n* + *const* for each *n*th leg.

**FIG. 4.**
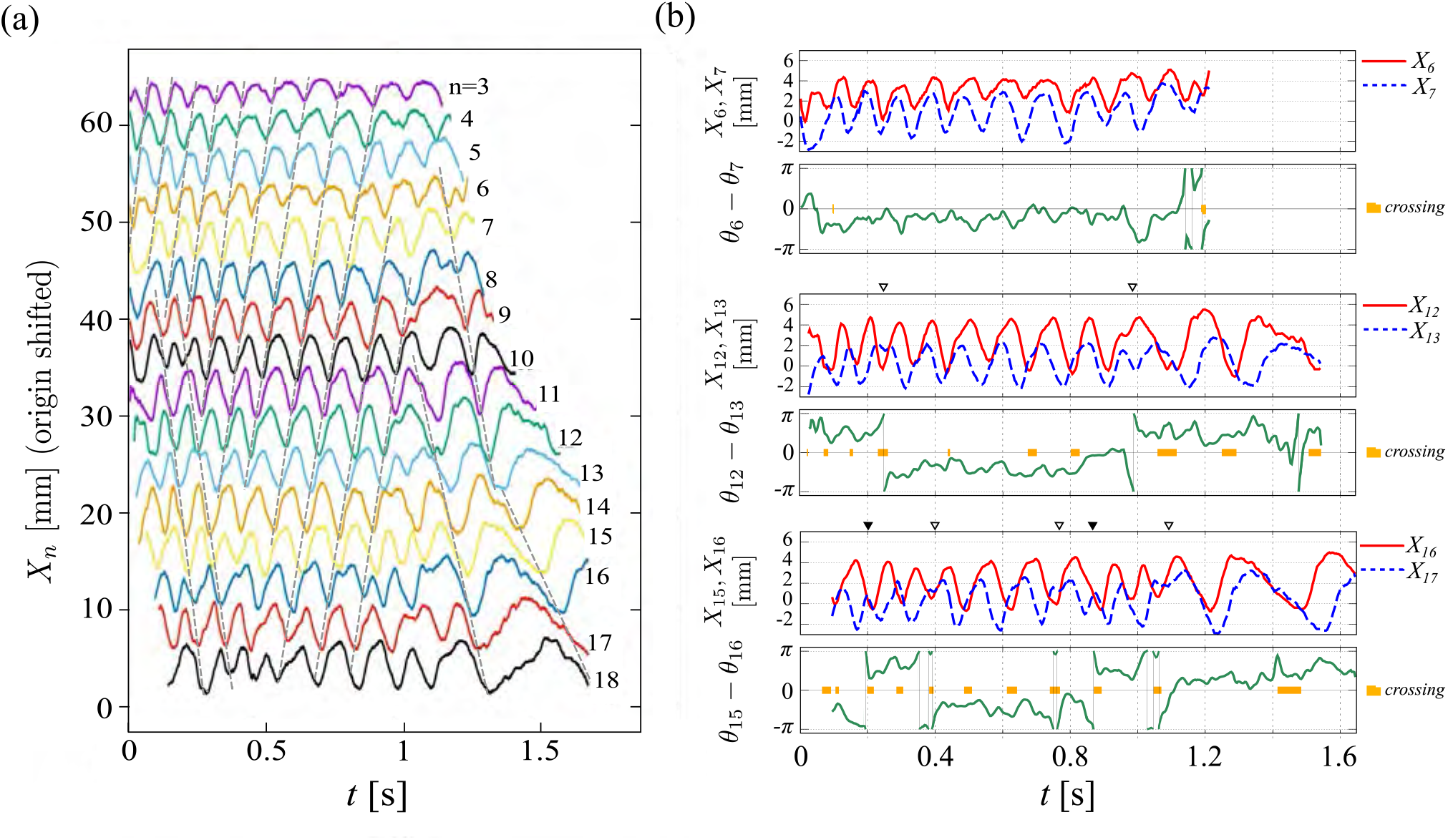
(a) Time-course of the leg-tip positions *X_n_*. The origin of the coordinate was shifted by (2*n* + const.) mm for each *n*. Wave fronts of direct and retrograde phase waves are shown by dotted lines. (b) Leg-tip positions of two adjacent legs (*X_n_*, *X*_*n*+1_) and their phase difference (*θ*_*n*_ – *θ*_*n*+1_) for *n* = 6,12, and 16. The time windows during which the two legs were crossed (i.e. *X_n_* < *X*_*n*+1_) are shown by thick orange lines in the phase difference plot. The inverted triangles indicate the places where the events mentioned in the text occurred.

In agreement with the phase plot in Fig. 3(a), direct waves were dominant in the anterior part of the centipede, while retrograde waves were found in the early and late stages in the posterior part of the centipede. Gait transition took place at the junction points of these two types of wavefronts. The waveforms of the leg-tip positions showed irregular patterns near the junction points. For example, the distance between adjacent legs was small at some of the junction points. Also, the waveform was split to have two peaks per cycle. The tip positions of two adjacent legs *X_n_* and *X*_*n*+1_ along with their phase difference Δ*θ_n_* = *θ_n_* — *θ*_*n*+1_ were plotted in Fig. 4(b) for the pairs *n* = 6,12, and 16, which represent legs in the anterior, middle, and posterior part of the centipede, respectively. The legs are considered to be crossed when *X_n_* < *X*_*n*+1_, and the time windows that satisfy this condition are plotted in thick lines in the phase difference plot. The phase difference Δ*θ_n_* was positive or negative when the two legs constituted local direct or retrograde waves, respectively. The transition between the direct and retrograde waves occurred when Δ*θ_n_* crossed either 0 or ±*π*. When it crossed ±*π*, one of the legs was going a cycle faster than the other. This is called a phase slip.

In the anterior part of the centipede (*n* = 6), the legs rarely crossed and were part of the direct waves for most of the time. Gait transitions with phase slips took place only at the last stage, and one of them (from retrograde to direct) was accompanied by a leg crossing. In the middle part of the centipede (*n* = 12), we observed transitions from retrograde to direct and back to retrograde. At the last stage (*t* > 1.45 s), we observed an irregular waveform with two more quick transitions that we regarded as an outlier. In both of the two main transitions (at *t* ≈ 0.25,0.99 s) the waveform of the back leg (*X*_13_) was disturbed with its peak split into two small peaks. One of the phase slips from local retrograde to direct waves was accompanied by a leg crossing. In the posterior part of the centipede (*n* = 16), we observed more frequent transitions between direct and retrograde waves. Leg crossings were observed in most of the cycles including those exhibiting transitions between retrograde and direct waves. The waveform of the back leg (*X*_17_) had a split peak at two of the phase slips (at *t* ≈ 0.4,0.75,1.1 s). The leg crossings observed with the retrograde-to-direct transition generally took place when the front (*n*-th) leg was in the retraction period (*dX_n_*/*dt* < 0) and the back ((n + 1)-th) leg was near the end of its protraction period (*dX*_*n*+1_/*dt* > 0). Therefore, the disturbance of the waveform of the back leg suggests that it was easier for the leg in the protraction period to fluctuate for a while. This is a natural way to adjust the phase difference, because the leg in the retraction period is touching the ground and thus its velocity in the body frame is strongly coupled to the body velocity. The reason why we found more leg crossings in the posterior part of the body might be attributed to the fact that the posterior legs are longer than the anterior legs. In the direct-to-retrograde transitions (at *t* ≈ 0.2, 0.86 s), the temporal increase of rotation frequency of the back leg by smaller power and/or recovery strokes seemed to provide the reversal of the phase relationship between successive legs.

### D. Continual gait transition (2): Movement on a slippery slope

An example of spontaneous transition between waves was observed when we put the centipedes on a slope with a slippery surface. We adjusted the angle of the slope so that the centipede could not climb up and slide off. In this situation, centipedes alternated running and resting at the same place, gaining or losing only several millimetres of ground. Figure 3b shows a typical gait transition while running after a short break. When the velocity was negative (*t* = 0 s), a source of locomotory waves appeared at the front leg, and then the wave propagated from head to tail and the posterior source disappeared (*I_w_* ↘ 0). After that, the global anterior source moved posteriorly (*I_w_* =0 ↗ 0.5) and remained around the middle for a short time (*t* ≈ 4 s). The source then again moved posteriorly toward the tail until it reached the hindmost leg (*I_w_* = 0.5 ↗ 1). This was accompanied by a large increase in dominant wavelength and decreased duty factor. The postero-anterior wave continued for a short time (*t* < 10 s). During this period, the dominant wavelength remained high (λ ≈ 6) and the duty factor remained low (*D* < 0.5). The velocity then increased again with the appearance of an anterior source (*I_w_* = 1 ↘ 0.5) and similar gait transitions were repeated. Over the entire experiment, the velocity variations were provided by variations in the span and the duty factor, while variations in the rotation frequency were very small.

### E. Switching surface condition: Transition from wet paper to plastic surface

To observe the influence of a sudden change in surface conditions, we covered half of the lane on the plastic plate with thin, wet paper. Figure 3c shows the time course of the gait transition that occurred at the surface condition boundary. We first induced a dash with a direct wave by tapping the centipede’s tail. When it approached the boundary and its antenna touched the plastic surface (*t* ≈ 0.5 s), it rapidly slowed down for a short time, then started crawling at a slower speed (*t* ≈ 0.75 s). After several anterior legs (pairs 1–8) passed the boundary with the same phase relationship among them, the 9th leg became a source leg, where each posterior leg approaching the boundary extended the period of the recovery stroke so that its footprint was identical with the anterior leg. As the centipede proceeded on the plastic surface, a global splitting wave appeared (*I_w_* = 1 ↘ 0.4) and the phase relationships were retained until all legs passed the wet-paper surface. This was accompanied by a large decrease of the dominant wavelength. After all legs had passed the wet-paper surface, the source leg again gradually shifted posteriorly (*I_w_* = 0.4 0.6).

When the centipede was on the wet paper, the phase difference between opposite legs was near *π*, which was observed in the swimming and dashing-to-slowdown conditions. However, after it moved onto the plastic surface, the phase differences were diminished in the legs that created local retrograde waves. It should be noted that the minimal phase difference during walking on the uniform surface was around *π*/4.

### F. Splitting waves as stable gaits

Splitting waves with a source leg in the anterior or middle part of the body was often observed in transient and stable gaits in various situations except swimming.

### G. Degree of body-support by legs during locomotion

To determine whether the body of the centipede was in contact with the substrate, tracks of centipedes walking or running over smoked paper were examined. During slow walking with low wave-indices, only traces of footprints were observed indicating that the body was fully supported off the ground by the legs. During fast running with large wave-indices, the tummy dragged along the track indicating that the body was not fully supported by the legs. This result is compatible with a previous report that used other Scolopendromorpha centipedes [17].

### H. Common features in gait transitions in various situations

The gait pattern was estimated by the wave-index *I_w_*, which is a discretised negatively correlated phase-order version of the mean wavenumber *κ_m_* (Fig. 5b). Figure 5a shows the gait pattern dependencies of the standard speed-control-related characteristics (rotation frequency, stride, span, and duty factor) across the five experimental conditions. The rotation frequencies were nearly constant, so the wave-index seemed to be independent of the rotation frequency. However, the stride had a clear positive correlation with the wave-index. In the small wave-index, the duty factor mainly contributed to this stride lengthening. In the middle and large wave-index, both the span and duty factor contributed to the stride lengthening. The dominant wavenumber was also negatively correlated with the wave-index (Fig. 5c). Fig. 5d shows the gait pattern dependency of the averaged dominant wavenumbers restricted in the local retrograde waves and local direct waves over the five conditions. In the middle and large wave-index, the decreased wavenumber in the local direct waves contributed to the further decrease of the mean and dominant wavenumbers. The wave-index and the speed did not have any global functional relationship Fig. 5e), but they seemed to have continuous relationships within each condition.

**FIG. 5.**
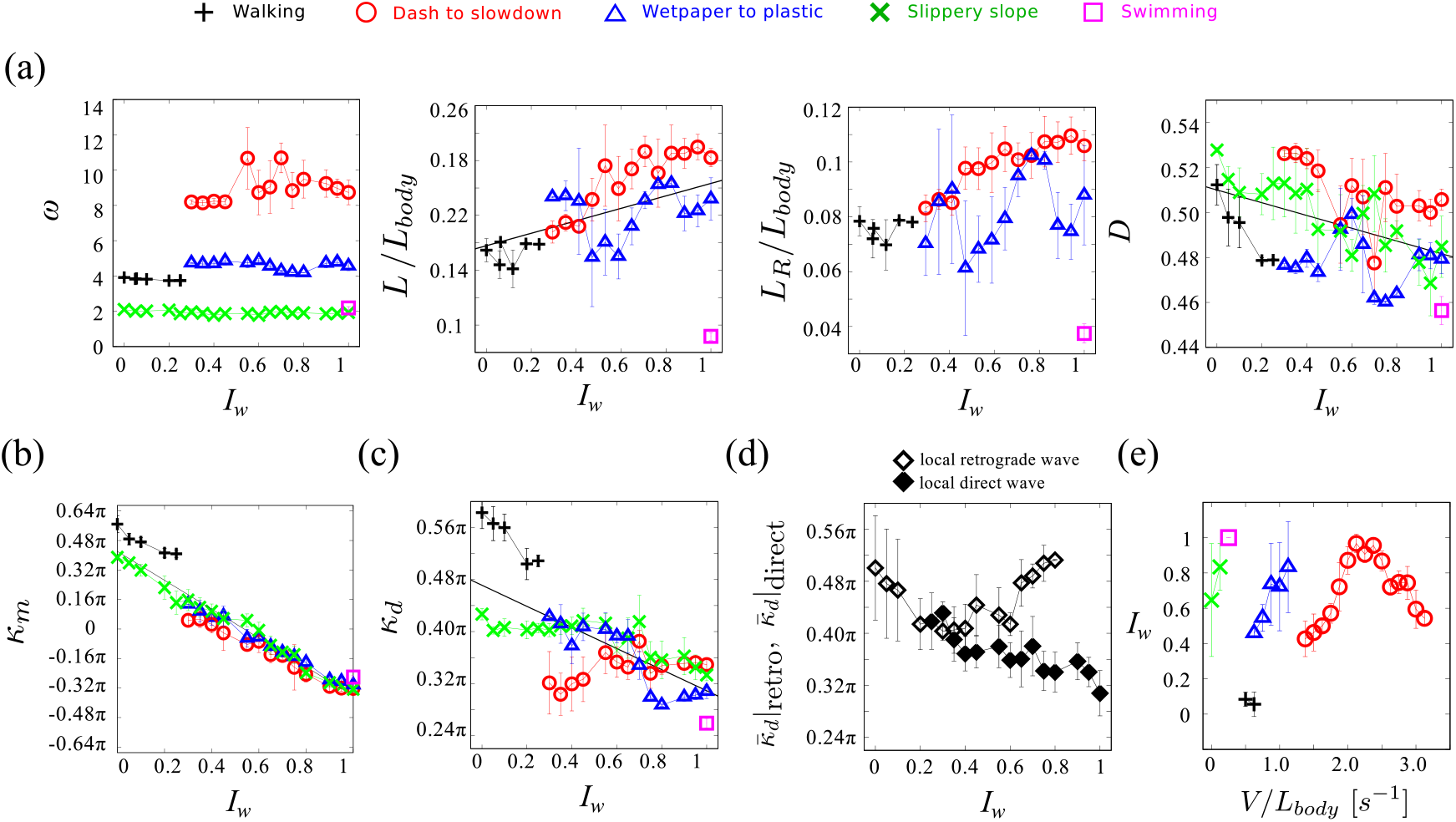
Relationships between wave-index and other leg-motion-related characteristics during walking, dash to slow-down, wet paper to plastic, slippery slope, and water crawling, which are shown by a plus sign (black), circle (red), triangle (blue), cross (green) and square (magenta), respectively. (a) Wave-index (*I_w_*) vs. rotation frequency (*ω*), relative stride (*L*/*L_body_*), relative span (*L_R_*/*L_body_*), and duty factor (*D*). In stride and pace graphs, the slippery slope case was excluded. The stride and duty factor data were fit to the lines *L* = 0.045*I_w_* + 0.16 (P = 0.010) and *D* = –0.028*I_w_* + 0.51 (P < 1e^−3^), respectively. These are shown by black solid lines. (b) Correspondence between wave-index (*I_w_*) and mean wavenumber (*κ_m_*). The dominant wavenumber data were fit to the line *κ_d_* = –2.451*I_w_* + 1.32*π* (P < 1.0e^−13^). (c) Wave-index (*I_w_*) vs. dominant wavenumber (*κ_d_*). The dominant wavenumber data were fit to the line *κ_d_* = —0.51*I_w_* + 0.47*π* (P < 1.0e^−8^). (d) Wave-index (*I_w_*) vs. averaged dominant wavenumber of local retrograde and direct waves for all experimental conditions 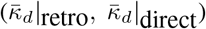. (e) Relative speed (speed divided by body length (*V*/*L_body_*)) vs. wave-index (*I_w_*).

## IV. DISCUSSION

### A. Gait transition in the centipede compatible with observed stride control and leg-crossing

The gait transitions from retrograde to direct locomotory waves were accompanied by stride increases that were provided by decreased duty factor over the entire wave-index, a constant span in the small wave-index, and increased span in the middle and large wave-index (Fig. 5b). The rotation frequency was independent of the gait transition. Moreover, the transitions were often preceded by the interference between successive legs (Fig. 4). To understand kinematic compatibility of these observations, it is convenient to consider a simple model where successive legs are treated as two identical rigid rods, and each rod swings around the perpendicular position (Fig. 6a). In this simple model, leg configurations are specified by a pair of angles (*γ_n_*, *γ*_*n*+1_). The trajectories corresponding to local retrograde waves and local direct waves are depicted by the counter-clockwise and clockwise closed quadrilaterals, respectively (Fig. 6b,c). Crossing of successive legs happens in the lower-right shadowed region (called the *leg-crossing region*), which is bounded by concave curves so that the trajectory with the smaller wavenumber is allowed to have the larger possible maximal span (Fig. 6b,c; see also Section IV D). The leg-crossing region depends on the ratio *R* = *d*/*l* where *ℓ* is the length of legs and *d* is the distance between the pivots of successive legs. The smaller *R* gives the larger area of crossing region. In the *S. rubiginosus, R* was 0.47 ± 0.06 (mean± sd; n=3). In the retrograde gait, as shown by the broken line in Fig. 6b, the decreased duty factor causes the trajectory to approach the leg-crossing region under the constant span condition.

**FIG. 6.**
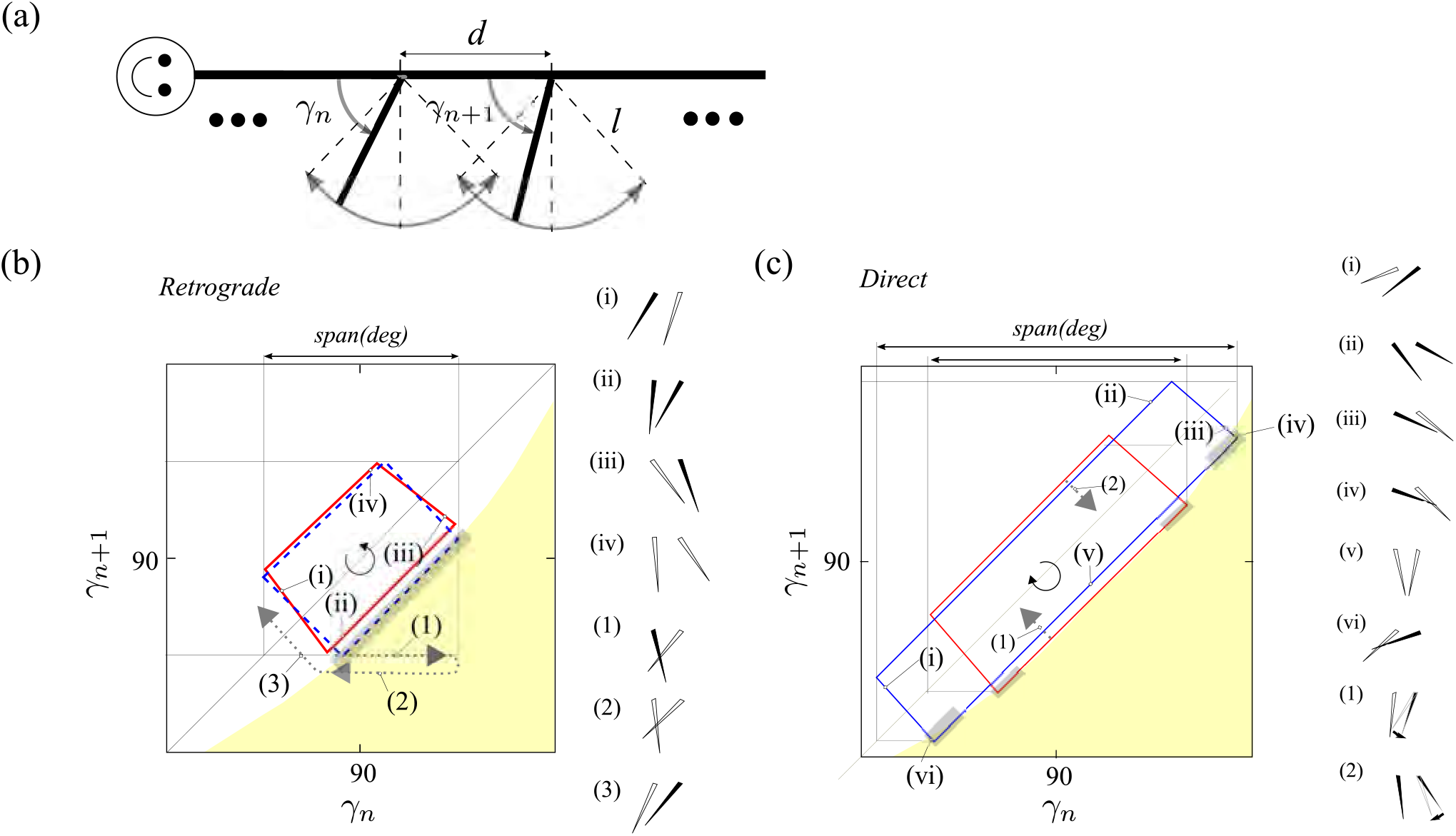
(a) Simple model for estimating the leg-crossing condition. The solid and blank legs show the retraction and protraction periods, respectively. For each ratio *R* = *d*/*ℓ*, the leg-touching conditions are expressed as concave curves in the *γ_n_* vs. *γ*_*n*+1_. (b) Effect of decreased duty factor on the leg-crossing in the retrograde gait. The shadowed region (yellow) shows the set of configurations of successive legs that cross each other (*R* = 2/3 case). The red solid-line trapezium shows the retrograde gait with a large duty factor (*D* > 0.5). Four successive legs arrangements on the trajectory are shown in (i)-(iv). The trajectory with the smaller duty factor (accompanied by a smaller wavenumber in the protraction period) with a constant span should approach the leg-crossing region. An impossible trajectory with duty factor *D* = 0.5 across the leg-crossing region is shown by the blue broken-line rectangle. The thin grey rectangle shadow indicates the regions where the leg-crossings occur. A schematic route of the local wave transition from retrograde to direct observed in Section IIIC (Fig.4) is denoted by the grey dotted-line and configurations of successive legs are shown in (1)-(3). (c) Effect of decreased wavenumber on the span increase in the direct gait. The thin grey rectangle shadows indicate the regions where the leg-crossings occur. Successive leg arrangements on the trajectory are shown in (i)-(vi). The two routes of local wave transition from direct to retrograde observed in Section IIIC (Fig.4) are denoted by grey dotted-lines and the successive legs arrangements are shown by (1) and (2), respectively.

One of the most remarkable differences between the retrograde waves and direct waves is in timing when the distance between the leg-tips of successive legs become smallest [9]. Because the dense part in the retrograde waves appears during the retraction period, the centipede may stumble if the trajectory intersects the leg-crossing region. This may give a strong motivator for the centipede to reverse the phase relationship between the legs. The grey dotted-line in Fig.6b shows a schematic route of the local wave transition from retrograde to direct observed in SectionIII C (Fig.4) where by the back leg staying at a certain small angle for almost one complete cycle, the phase relationship between the successive legs is inverted. In contrast, the dense part in the direct waves appears during the protraction period. Therefore, the crossing legs can avoid interfering with each other varying their elevation angles, as reported for a centipede such as *Scutigera* ([8] p.367). This enables the trajectory to partially intersect the leg-crossing region so that the possible maximal span in the direct waves is larger than that in the retrograde waves at each wavenumber. It should be noted that the trajectory in the protraction period of direct waves must get out of the leg-crossing region once because the back leg should be lower than the upper side of the front leg before beginning its retraction.

### B. Body-size dependency of locomotory waves

The body-size dependency of the gait pattern is an interesting problem. Although we examined only putative adult centipedes, it seemed that the smaller centipedes tended to use direct waves more than the larger ones. A comprehensive study of this phenomenon will be left for future studies.

### C. Comparison of gait transition with other scolopendromorpha

Among the five orders of the centipede class, geophilomorpha and scolopendromorpha have short legs and are slow, strong movers that usually use retrograde waves [8]. Manton reported the gait transition of *Scolopendromorpha scolopendra cingulata* in detail [18]. In the Scolopendromorpha centipede, which is similar to the *S. rubiginosus L. Koch* used in this paper, increased stride was strongly correlated with decreased wavenumber and duty factor, although the direction of locomotory waves was in the same direction. Its wavelength reached up to 13 [#leg], which is twice of that of *S. rubiginosus L. Koch*. In the gait with the larger wavelength, the lateral undulation became deeper in the *S. cingulata*. However, in *S. rubiginosus L. Koch*, except when it was swimming, the lateral undulation was not conspicuous, which is compatible with a previous study of other *Cryptops* [18]. We roughly estimated the ratio *R* from the photo of *S. cingulata* in [18]. Interestingly, the *R* is 0.58±0.04 (mean±sd; n=1) which is larger than that of *S. rubiginosus*. Further investigation of this comparative aspect will be left for future studies.

### D. Fluid mechanical viewpoint for direct waves in centipede swimming

We investigated the advantage of direct waves in limbed organism swimming. The Reynolds number of a swimming centipede was estimated to be on the order of 10, which is similar to that of young crayfish [19]. For example, Re = *UL*/*ν* = 74, where *U* = 2.64 cm/s is the mean maximal speed of limb movement during a power stroke, *L* = 2.77 cm is the mean length of the underwater portion of a limb (from femur to pretarsus), and *μ* is the kinematic viscosity of water (0.01 cm^2^/s at 20 °C). It is known that limb coordination of swimming crustaceans, including crayfish and shrimp, occurs as direct waves. A study using a computational fluid dynamics model [19] showed that direct waves were the most effective stroke pattern across the range of biologically relevant Reynolds numbers for swimming crustaceans (10 < Re < 1000). This is because the power stroke associated with the sparse phase of the direct wave encloses a larger volume of fluid and generates much more positive flux than other swimming patterns. This hydrodynamic effectiveness may also hold for centipedes. This is supported by the existence of the centipede-like crustacean ! *remipede*, which uses direct waves as its swimming gait [20]. In contrast, another report [21] showed that some centipedes without appendages (e.g., *Scolopendra cataracta)* swim in an eel-like way using vigorous horizontal undulation waves that propagate antero-posteriorly. Further investigation of such differences among centipedes using water crawling patterns is left as future work.

## AUTHORS’ CONTRIBUTIONS

SK and TN designed the study. SK conducted the experiments. SK and NU analysed the data and wrote the manuscript. All authors gave final approval for publication.

## ACKNOWLEDGEMENTS

SK acknowledges helpful comments on data interpretation from Dr. Helmut R. Brand (Department of Physics, University of Bayreuth). We thank Melissa Gibbons, PhD, from Edanz Group (www.edanzediting.com/ac) for editing a draft of this manuscript.

## Estimation of leg-crossing condition in the simple model

To illustrate the relationship among the interference of successive legs, duty factor, wavenumber, and possible maximum span, we used a simple model in which the legs move with a prescribed symmetric swinging motion around the perpendicular position, mimicking the power and recovery strokes of the legs (Fig. 7a). Specifically, the nth leg-tip position was prescribed to be (*x_n_*, *y_n_*) = ((*N* – *n*)*d*, 0) + *ℓ*(cos *γ*_*n*+1_, sin *γ*_*n*+1_) (*n* = 1, 2,…, *N*), where all legs are straight rigid rods with the same length, *ℓ*, and the distance between pivots of successive legs, *d*, was constant.

**FIG. 7.**
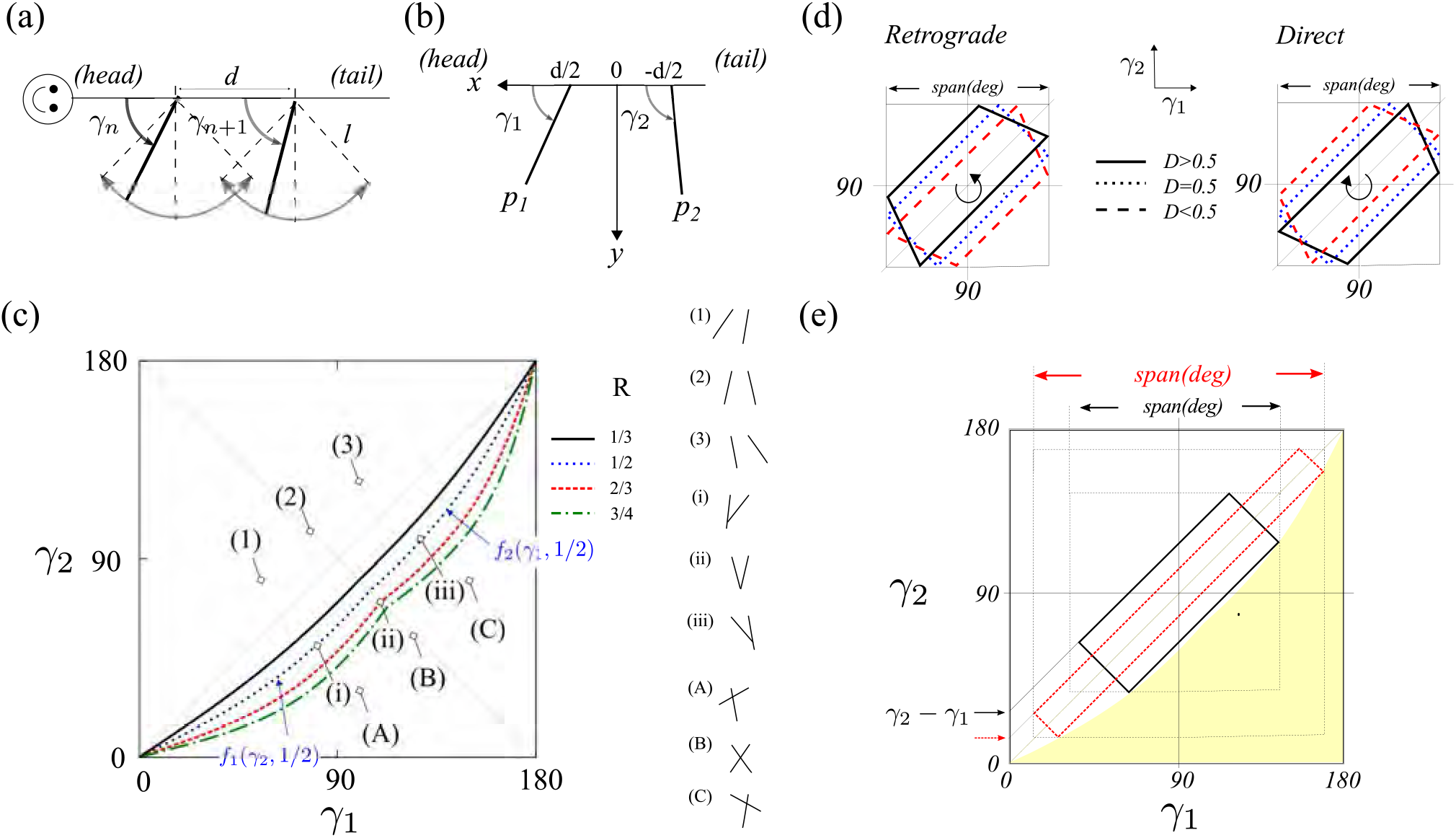
Simple model illustrating the leg-crossing condition for successive legs. (a) Prescribed symmetrical swing motion of each leg around the perpendicular position (top view). (b) Introduction of a specific (*x, y*)-coordinate system representing the leg-tip position to calculate the touching curves. (c) The touching curves of four different *R* on the (*γ*_1_, *γ*_2_)-plane. Configurations of successive legs are shown in (1)-(C). (d) Effect of decreased duty factor under constant span on the trajectory. Local retrograde waves and local direct waves are shown in the left and right panels, respectively. (e) Effect of decreased wavenumber under even duty factor (*D* = 0.5) and non-leg-crossing condition (*R* = 1/3) on the span. The wavenumber and span of the long-narrow red dotted-line rectangle are smaller and longer than those of the black short-wide solid-line rectangle).

Now, we consider when successive legs cross. Without any loss of generality, we considered the *N* = 2 case and set *P*_1_: (*x*_1_, *y*_1_) = (*d*/2, 0) + *ℓ*(cos *γ*_1_, sin *γ*_1_) and *P*_2_: (*x*_2_, *y*_2_) = (–*d*/2, 0) + *ℓ*(cos *γ*_2_, sin *γ*_2_) (Fig. 7b). Then, the necessary condition for the leg crossing point *P*_3_ was *x*_3_ = *d*(tan *γ*_1_ + tan *γ*_2_)/2(tan *γ*_1_ – tan *γ*_2_). The touching of aleg-tip to another leg occurs when the 2nd leg-tip touches the 1st leg and vice versa, which satisfies the conditions |*P*_2_*P*_3_| = *ℓ* and |*P*_1_*P*_3_| = *ℓ*, respectively. From a simple calculation under *R* = *d*/*l*, the first case occurs when *x*_2_ > 0 and 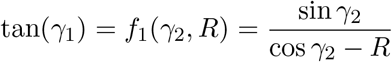. The second case occurs when *x*_1_ < 0 and 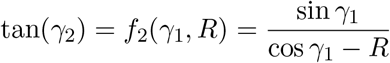.

The graphs of *f*_1_ and *f*_2_ in the (*γ*_1_, *γ*_2_)-plane are concave curves that meet on the *γ*_2_ + *γ*_1_ = 180 line. The curves are called the touching curves. The graphs of touching curves for *R* = 3/4, 2/3,1/2,1/3 are shown in Fig. 7c. Leg crossings occur in the lower right region below the touching curve. (A)-(C) in Fig. 7c are three examples in the crossing region. The smaller *R* gives the larger area of crossing region. The trajectories corresponding with local retrograde waves and local direct waves are depicted by the counterclockwise and clockwise closed quadrilaterals, respectively. The swing angular speeds were assumed to be common in all legs and were uniform during retraction and protraction periods, but their ratio (the duty factor) could be changed. It should be noted that the smaller duty factor provided a smaller wavenumber during the protraction period but a larger wavenumber during the retraction period. Fig. 7d illustrates the effect of a decreased duty factor under the same span on the trajectory’s shape. In the retrograde waves, the trajectory during the retraction period approached the leg crossing region, and in the direct waves, the trajectory moved away from the leg crossing region. Fig. 7e illustrates the effect of decreased wavenumber under the non-leg-crossing condition on the trajectory span. The possible maximum span increased as the wavenumber decreased.

## List of symbols

*t*: time [s]
*γ_n_*: swing angle of the *n*th leg [deg]
*θ_n_*: phase of the *n*th leg [rad]
*T_R_*: retraction period [s]
*T*: pace duration [s]
*L_R_*: span [mm]
*L*: stride [mm]
*D*: duty factor
*ω*: rotation frequency [s^−1^]
*I_ω_*: wave-index
*λ_d_*: dominant wavelength [#leg]
*κ_d_*: dominant wavenumber [rad/#leg]
*κ_m_*: mean (signed) wavenumber [rad/#leg]
*X_n_*: putative leg-tip position along the body axis [mm]
*b_n_*: coordinate of the base of the *n*th leg [mm]
*r_n_*: length of the *n*th leg [mm]

